# Direct measurements of mRNA translation kinetics in living cells

**DOI:** 10.1101/2020.10.12.335505

**Authors:** Mikhail Metelev, Ivan L. Volkov, Erik Lundin, Arvid H. Gynnå, Johan Elf, Magnus Johansson

## Abstract

Ribosome mediated mRNA translation is central to life as we know it. The cycle of translation has, however, not been characterized in a living cell. Here we have developed a live-cell ribosome-labeling method, which allows us to characterize the whole processes of finding an mRNA and translating it, using single-molecule tracking techniques. We find that more than 90% of both bacterial ribosomal subunits are engaged in elongation at any particular time, and that neither of the subunits, in general, continues translation from one open reading frame to the next on a poly-cistronic mRNA. Furthermore, we find that a variety of previously published orthogonal ribosomes, with altered anti-Shine-Dalgarno sequences, show significant binding to endogenous mRNAs, with the rate of translation initiation only modestly affected. Hence, our results suggest that other mRNA elements than the SD sequence play major roles in directing the ribosome to the correct translation start sites.

## INTRODUCTION

Translation of mRNAs into proteins is one of the most fundamental molecular processes in all living organisms. Central to this process is the ribosome, a highly conserved macromolecular machine consisting of two subunits, the small (30S in bacteria), and the large (50S in bacteria). mRNA translation has been extensively studied over the decades, and a wide variety of bulk biochemical approaches and structural techniques have helped painting a detailed picture of the process^1^. Recent advances in cryogenic electron microscopy continue to provide new snapshots of ribosomes with numerous ligands, capturing multiple conformations, and revealing hidden structural dynamics^2–3^. Emerging single-molecule techniques have, at the same time, helped to connect the conformational dynamics of the ribosome to transient interactions with ligands^4^. Until now, however, most studies of translation kinetics have been performed using reconstituted *in vitro* systems. Although extremely powerful, studies of an isolated system can never fully account for the complexity emerging from inter-connected molecular processes occurring in the crowded environment of the living cell. The development of new methods that allow direct measurements of mRNA translation kinetics inside the cell is, hence, crucial for a complete understanding of mRNA translation and its regulation.

*In vivo* super-resolution based single-molecule tracking techniques, have been successfully used to differentiate functional binding states of key macromolecules (recently reviewed in e.g.^5^). Initial applications of this approach has mostly been limited to the identification of cellular localizations of molecules in different functional states, whereas the dynamics of the processes has been out of reach due to limited photostability of the fluorescent proteins which have dominated the field of fluorescence microscopy. The development of new approaches to label biomolecules of interest, exploiting the brightness and photostability of small organic dyes, has opened up new possibilities for live-cell kinetics studies. Such approaches include, e.g., tracking of single dye-labeled tRNAs for translation kinetics measurements^6^, and measurements of spatiotemporal dynamics and translational activity of individual mRNAs^7–11^. The latter was accomplished by introducing arrays of labeling sites on both mRNA and nascent peptides, hence achieving multiple-fluorophore labeling of active translation sites.

In the present work, we introduce a system to directly study *in vivo* kinetics of translation initiation and elongation. We demonstrate that we can readily detect subtle changes in translation initiation kinetics as a response to antibiotics treatment or due to modifications in the ribosome itself. Our results further shed light on long-standing questions regarding how the bacterial ribosome finds the correct translation start site, and how polycistronic mRNAs are translated in the bacterial cell.

## RESULTS

### Construction of strains for ribosome labeling

To track 30S and 50S ribosomal subunits in live *Escherichia coli* we constructed strains in which the gene for HaloTag^12^ was inserted at the C-terminus of the chromosomal locus for ribosomal proteins L1, L9, L19, L25, or S2 (Fig. 1A). Strains expressing L9-HaloTag and S2-HaloTag showed minor negative effect on cell growth, cells in which L25 was mutated showed slower growth, while strains expressing L19-HaloTag or L1-HaloTag showed pronounced growth defects (Fig. S1A).

**Fig. 1.**
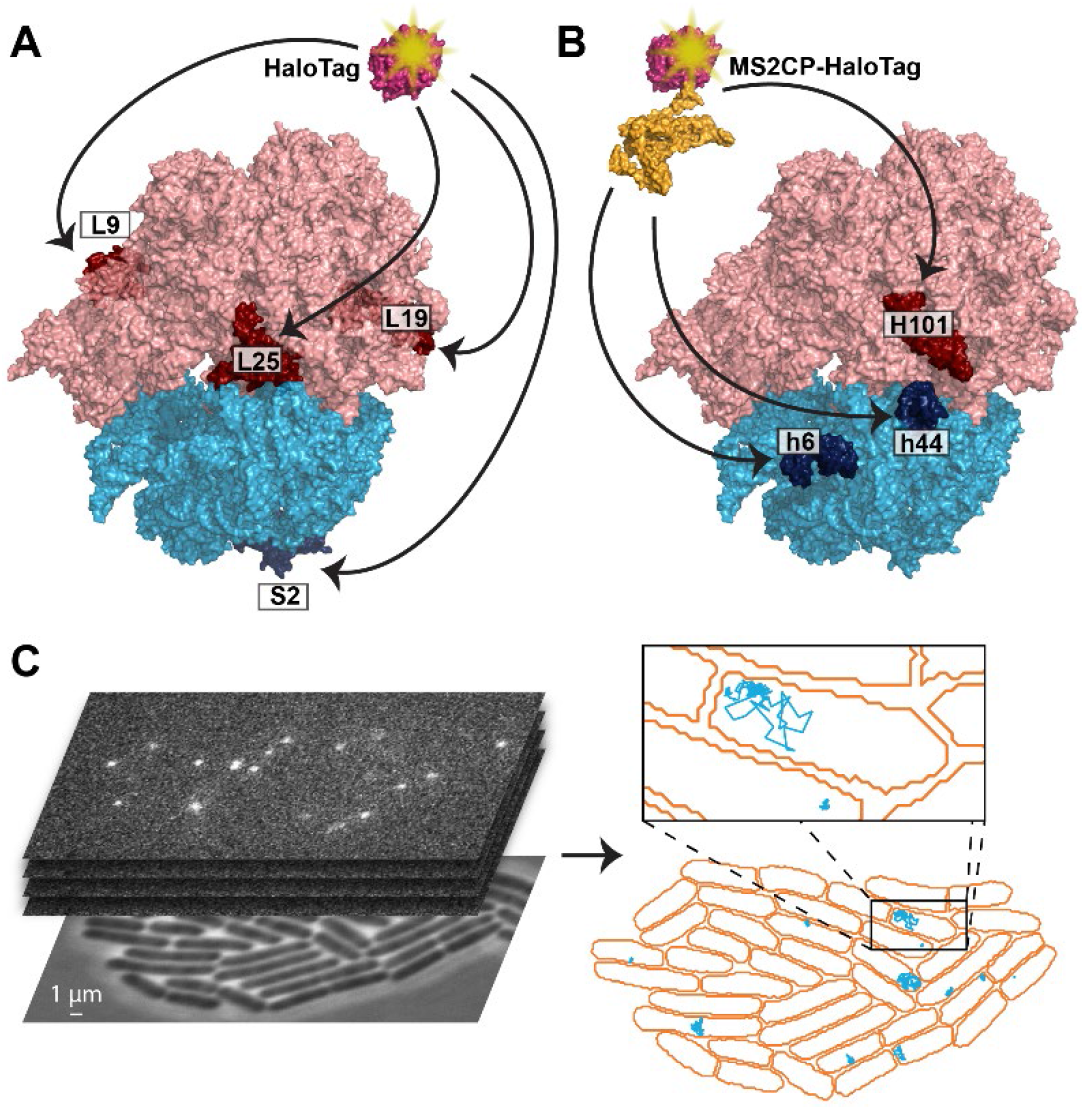
Single-molecule tracking of ribosomal subunits. **A.** Site-specific labeling of ribosomal subunits was achieved by genetically fusing the HaloTag protein to a ribosomal protein (L9, L19, or L25 of the 50S subunit, and S2 of the 30S subunit). **B.** Site-specific labelling of only a subpopulation of ribosomal subunits was achieved by introducing the MS2 RNA aptamer into a surface exposed helix of ribosomal RNA (23S rRNA H101, 16S rRNA h6 or h44), to which a co-expressed MS2CP-HaloTag fusion protein binds. **C.** Fluorescence time-lapse movies of cells were acquired, and diffusion trajectories of labeled ribosomal subunits were automatically detected in cells segmented based on phase-contrast images.

Labeling of ribosomes by means of ribosomal proteins fused to the HaloTag provides a tool for tracking of the total pool of ribosomal subunits present in cells. However, this approach lacks discriminative power when only a subpopulation of ribosomes with specific properties is of interest. In several studies it has been shown that mutated ribosomes with desirable modifications can be isolated by introducing an MS2 coat protein (MS2CP)^13–15^. Hence, to be able to track only a subpopulation of ribosomes, we inserted the MS2 RNA aptamer into the rRNA, which is then used of ribosomes, we introduced the different plasmids with the modified *rrnB* operons into the *E. coli* SQ171 strain, which lacks all chromosomal rRNA operons. We found that extension of h6 of the 16S rRNA did not affect the growth rate of the cells, while cells with modified 16S rRNA h44 and 23S rRNA H101 showed significant growth defects (Fig. S1B).

### Tracking of single fluorescent 30S and 50S subunits in live cells

For single-molecule tracking experiments we selected the L9-HaloTag and S2-HaloTag strains, as well as a strain containing both the plasmid for expression of h6-modified 16S rRNA and a plasmid encoding the MS2CP-HaloTag fusion. To fluorescence label the molecules fused to the HaloTag, cells in exponential growth phase were incubated with the JF549 HaloTag ligand which penetrates the cells and rapidly and specifically forms a covalent bond with the HaloTag protein^16^. After extensive washing, the cells were sparsely spread on an agarose pad, grown to mini colonies, and imaged at 37 °C (Fig. 1C). For all strains tested, we observe that practically all (~99%) cells continued growth on the agarose pad and contained fluorescently labeled molecules. Labeled molecules were stable for several hours and were homogeneously distributed within mini colonies produced from each single labeled cell over time as a result of cell division (Fig. S2). Mini colonies were imaged using stroboscopic laser illumination with 3 ms laser pulses per 30 ms camera exposure frame. The resulting microscopy data obtained from more than 100 individual colonies were processed using a previously developed pipeline for automatic cell segmentation, fluorescent molecules detection, and building of single-molecule diffusion trajectories6 (Fig. 1C). The apparent average diffusion coefficient for the HaloTag molecules was very similar in all the selected strains (Fig. S3A) and consistent with the expected diffusion for ribosomal particles^17–18^, i.e. around 0.1 μm^2^/s. As a control, we also tracked free labeled HaloTag proteins in the genetic background strain, as well as MS2CP-HaloTag in cells expressing the *rrnB* operon without insertion of the MS2 aptamer. In both these cases, the apparent average diffusion coefficient of the labeled molecules was approximately two orders of magnitude higher (Fig. S3B), matching what was previously reported for freely diffusing fluorescent proteins^19^. Hence, the HaloTag fusion to ribosomal proteins allow specific labeling of ribosomal subunits, whereas the presence of the MS2 aptamer in 16S rRNA allows specific labeling of a subpopulation of 30S for *in vivo* tracking.

### HMM analysis of diffusion trajectories

The high brightness and stability of the JF549 dye allowed us to track ribosomal subunits up to hundreds of frames (Fig. S4), which lead to the visual observation that the ribosomal particles do not diffuse homogeneously over time, but rather persist in different long-lived relatively discrete diffusional states (Supplementary Movie S1).

To obtain quantitative information on diffusion states and the transitions between these, we analyzed the diffusion trajectories using a Hidden Markov Modeling (HMM) approach^20^ previously used to extract tRNA-ribosome binding kinetics^6^. In this approach, all trajectories are fitted to a pre-set number of diffusion states, using the respective diffusion coefficients (*D*) and the transition frequencies between these states as fitting parameters. Based on the results from previous single-molecule tracking of ribosomal subunits labeled with fluorescent proteins^17–18^, we expected that the HMM analysis should be able to distinguish freely diffusing ribosomal subunits from translating subunits bound to mRNA. However, we also anticipated that the ‘bound’ state, in particular, might not be well defined, but rather consist of a continuum of diffusion states representing everything from a comparably rapidly diffusing single 30S or 70S bound to an mRNA, to slowly diffusing polysomes, perhaps also tethered to the chromosome or bound to translocons in the cell membrane^17, 21–22^. Hence, the diffusion trajectories were fitted to models with 2-8 discrete diffusion states. The average diffusion coefficient distributions are very similar between independent experiments (Fig. S5), and for different experimental replicas we observe good overall reproducibility in the diffusion states predicted by HMM (Tables S1-S3). To achieve convergence in the kinetics data, however, the results for all experiments below are presented from cumulated data from several independent experiments (2-5), with standard errors estimated by bootstrapping in the HMM fitting procedure.

For both the 30S and 50S tracking data, independent of model size, we find the largest fraction of ribosomal subunits diffusing at 0.03-0.06 μm^2^/s, and a smaller fraction at around 0.4 μm^2^/s (Fig. S6, Tables S4-S6). In models with more HMM states, the slower state is divided into several states, in agreement with our expectation of a loosely defined ‘bound’ state. Further, with model sizes of 6 or more states, low populated states (< 0.5%) at unphysiological ribosome diffusion rates > 3 μm^2^/s appear. Based on our previous analysis of simulated fluorescence microscopy6, we tentatively assign these very rapid diffusion states as tracking artefacts, possibly in combination with a very small fraction of dissociated HaloTag protein.

### Diffusion of free ribosomal subunits

Efficient initiation of translation for the majority of the *E. coli* genes depends on the presence of the Shine-Dalgarno (SD) motif, with a consensus sequence GGAGG located upstream of the AUG start codon. The SD motif facilitates recruitment of the 30S subunit via RNA-RNA base-pairing with the corresponding CCUCC motif at the 3’ terminus of 16S rRNA, referred to as the anti-Shine-Dalgarno sequence (ASD)^23^. Alternations in the ASD have been shown to render 30S subunits unable to efficiently initiate translation of endogenous mRNAs, while mRNAs with the corresponding modifications in the SD motif were specifically translated only by such mutated, orthogonal, subunits^24–25^. Hence, to specifically study the diffusion of unbound 30S subunits, we introduced the MS2 RNA aptamer into h6 of the orthogonal 30S (O-30S) developed in the Chin lab25, and co-expressed these with the MS2CP-HaloTag fusion.

We noticed that expression of O-30S subunits was moderately toxic for the *E. coli* cells, and that a C1400U mutation in the 16S rRNA of the O-30S appeared frequently, which alleviates this toxicity. The C1400 residue in 16S rRNS residue lies at the center of the decoding site of 16S rRNA, and other mutations of C1400 in ASD-altered ribosomes greatly impact the overall activity of the ribosomes for both endogenous mRNAs and orthogonal mRNAs^26^. Hence, it is likely that O-30S subunits are still significantly involved in translation of endogenous genes, perturbing the cell proteome, and that the C1400U mutation reduces this activity^27–28^.

We tracked both O-30S and O-30S-U1400 and compared their diffusion with that of normal 30S subunits. The average diffusion of O-30S and O-30S-U1400 was found to be 0.4 μm^2^/s and 0.5 μm^2^/s, respectively (Fig. 2A). This is significantly higher than 0.1 μm^2^/s observed for 30S subunits with unaltered ASD (Fig. 2A), thus suggesting that, in deed, both types of O-30S bind mRNA to less extent. From the HMM analysis of O-30S trajectories we find that, for all model sizes, the occupancy (steady-state fraction) in the fast (0.25-0.70 μm^2^/s) diffusion states increases dramatically, to ~50% and ~80% for O-30S and O-30S-U1400, respectively, compared to < 10 % for ribosomes with unaltered ASD (Fig. 2B and Tables S6-S8). Hence, a threshold for diffusion of 30S subunits at 0.25 μm^2^/s would enable us to separate freely diffusing 30S from 30S participating in translation. The difference in diffusion for O-30S and O-30S-U1400, further suggests that O-30S subunits partially retain their ability to bind and translate endogenous mRNAs, in agreement with results of ribosomal profiling for orthogonal ribosomes^28^, and that mutation of U1400 dramatically reduces their affinity to mRNA.

**Fig. 2.**
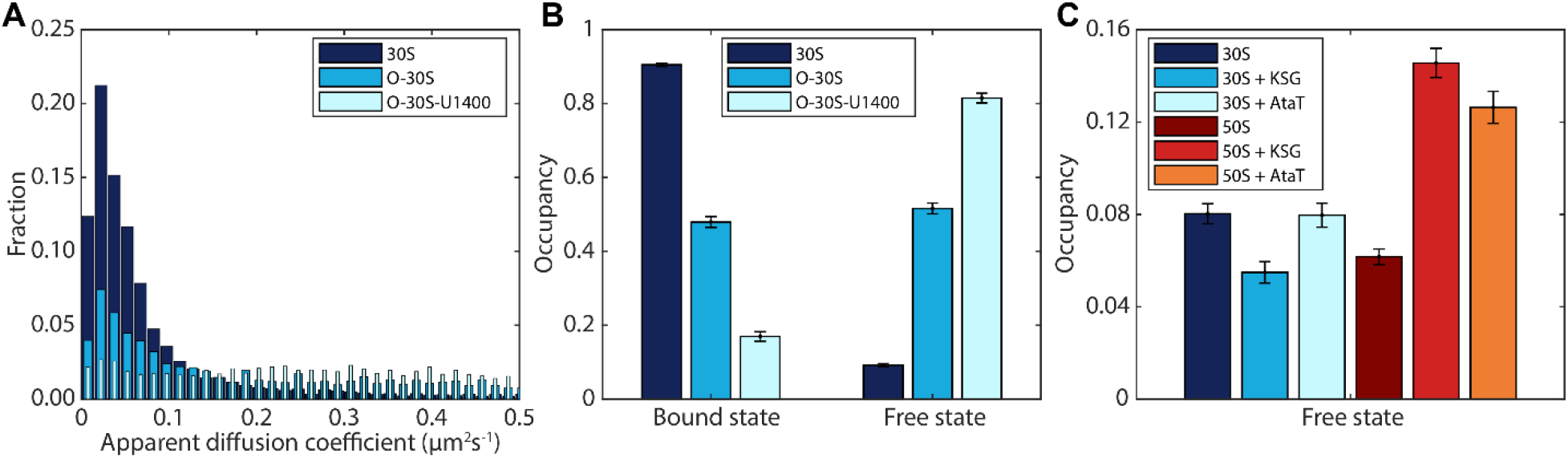
Distinction between freely diffusing and mRNA-bound ribosomal subunits **A.** Distribution of apparent diffusion coefficient, estimated from mean-squared-displacement analysis of diffusion trajectory segments of 7 frames. **B.** HMM-estimated occupancy (steady-state fraction) of 30S subunits in mRNA-bound or freely diffusing state. **C.** HMM-estimated occupancy of 30S and 50S subunits in the freely diffusing state in presence or absence of 50 μg/ml KSG or the AtaT toxin. Panels B and C show coarse-grained HMM results from a 6-state model, using 0.25 μm^2^/s as cutoff between bound and free subunits. The complete results for all model sizes (2-8) are shown in Tables S6-S8. Error bars represent bootstrap estimates of standard errors.

To further validate the threshold, and to investigate whether the same threshold can be used for 50S subunits, we tracked 30S and 50S in cells treated by kasugamycin (KSG), an aminoglycoside antibiotic that specifically target translation initiation^29^. It has been shown *in vitro* that KSG moderately affects the binding of 30S subunit to an mRNA, but strongly inhibits its affinity to initiator fMet-tRNA^fMet^ ^30^. Thus, KSG should prevent the formation of the 70S initiation complex, and, in particular, the arrival of 50S is expected to be delayed. *E. coli* cells containing either L9-HaloTag or S2-HaloTag were grown on an agarose pad for 2 hours, after which the pads were soaked in media containing 50 μg/ml KSG (0.2 times MIC, at which cells show 1.5 times longer doubling times). From the HMM analysis of diffusion trajectories in cells subjected to KSG treatment, we observe that the occupancy in diffusion states in the interval of 0.25-0.70 μm^2^/s (‘Free state’) increases from 6% to 15% for 50S, whereas the free state occupancy of 30S instead decrease slightly (from 8 % to 6%, Fig. 2C, Tables S9-S10). The difference in proportion of free 30S and 50S in KSG treated cells, compared to the equal proportion in untreated cells (Fig. 2C), hence shows that our analysis is capable of distinguishing free 30S subunits from 30S subunits bound to an mRNA awaiting 50S joining. When cells were treated with higher KSG concentration (500 μg/ml, 2X MIC), the diffusion distribution for both 30S and 50S particles changed radically, with ~50% occupancy in the freely diffusive state (Tables S11-S12).

To corroborate the result from KSG treatment even further, we performed a complementary experiment using the AtaT toxin. In this experiment, strains encoding S2-HaloTag and L9-HaloTag were transformed with a plasmid for low level of expression of AtaT. AtaT is a toxin from the recently characterized toxin-antitoxin complex, AtaRT, which specifically modifies Met-tRNA^fMet^ by acetylating the free amino group of the methionine31. This modification prevents the interaction between tRNA^fMet^ and IF2 which leads to inefficient formation of the 70S initiation complex^31^. Effectively, expression of AtaT toxin should decrease the concentration of available initiator fMet-tRNA^fMet^, slowing down the final steps of translation initiation, including 50S subunit joining.

The results from tracking 30S and 50S subunits in AtaT producing cells were, as expected, very similar to what was observed in cells treated with 50 μg/ml KSG, showing an increase in the proportion of free 50S subunits whereas no significant effect was found on the 30S subunits (Fig. 2C, Tables S13-S14). Importantly, the experiments with KSG and AtaT treatment show that once the 30S subunit established interactions with an initiation site, it is unlikely to leave it and instead awaits 50S arrival.

### Kinetics of translation elongation

In addition to the occupancy in each diffusion state, the HMM analysis also provides the estimated transition frequencies between these states, or conversely, the average dwell-time in each state. To study specifically the transitions between free subunits and mRNA-bound subunits, we coarse-grained the HMM models of each size obtained for 50S subunits (L9-HaloTag) and 30S subunits (S2-HaloTag and h6-MS2-HaloTag) into three states in accordance with the control experiments presented in the previous section: mRNA bound ribosomal subunits at *D* < 0.25 μm^2^/s, freely diffusing subunits with *D* in the interval of 0.25-1.5 μm^2^/s, and a third state at *D* > 1.5 μm^2^/s in which tracking artifacts detected in larger models appear. Both the occupancy and the dwell-time in the coarse-grained functional states converge well for both 30S and 50S subunits with increasing model size (Fig. 3A-B). For simplicity and clarity, in the following discussions, we thus present only coarse-grained numbers from the 6-state models. Whereas absolute numbers are slightly different between different model sizes, our conclusions do not depend on exact model size ≥ 4 states.

**Fig. 3.**
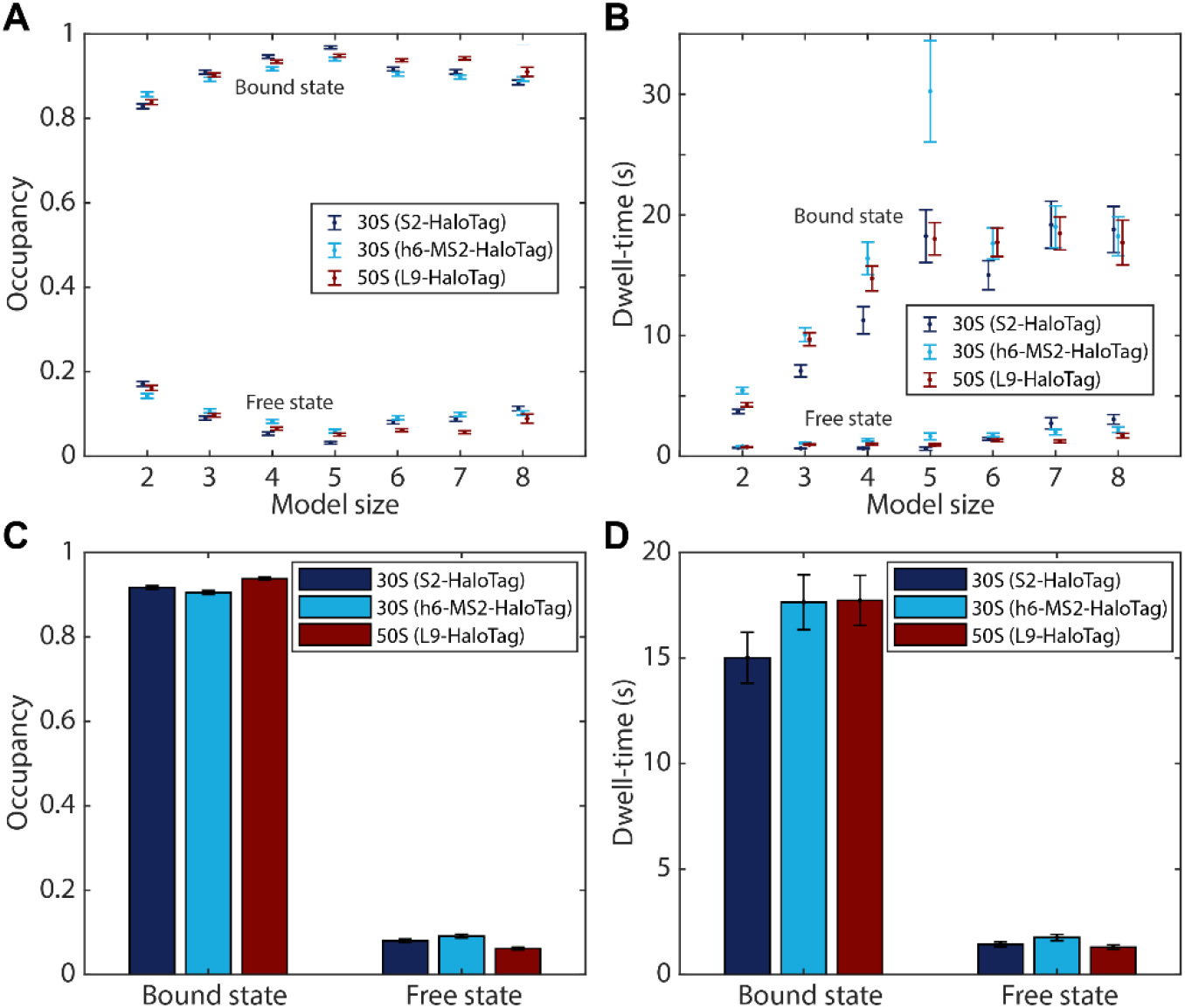
Translation kinetics from coarse-grained HMM-results. **A-B.** Estimated occupancy and dwell-time of ribosomal subunits in the mRNA-bound state or the freely diffusive state for different HMM fitting model sizes. Full models (Tables S4-S6) were coarse-grained into three states: mRNA-bound, free, and artifacts, using thresholds 0.25 μm^2^/s and 1.5 μm^2^/s. **C-D.** Bar plots of model size 6 data as in panels A-B. Error bars represent bootstrap estimates of standard errors.

To investigate the kinetics of translation inside rapidly growing *E. coli* cells, we first compare the steady-state fraction of 30S and 50S subunits in the free or mRNA-bound diffusion states. For the 50S subunit, we find that 94% are at any given moment engaged in translation. The corresponding value for the 30S subunit is 92% or 91% for S2 or h6 labeling respectively (Fig. 3C). Hence, the fraction of free subunits, 6-9%, is somewhat lower than what has previously been reported using other methods (i.e. 15%^32^. However, it has also been estimated that precursors of 30S and 50S subunits comprise approximately 10% of all subunit particles^33^. Since in our experiments, data were acquired 2 hours after labeling, and we see no or negligible exchange of the ribosomal proteins and MS2CP-HaloTag between different subunits, we most probably track only the diffusion of fully processed particles. Hence, from our analysis we estimate that ≥ 90% of the *mature* fraction of both subunits are actively translating at any given moment.

From the HMM estimated transition frequencies of the ribosomal subunits between the diffusion states, we find that the 50S subunits spend, on average, 18 +/− 1 s bound to an mRNA, whereas the time spent freely diffusing between binding events is on average 1.3 s +/− 0.1 s (Fig. 3D). For the 30S subunit, the corresponding numbers are 15 +/− 1 s and 1.4 s +/− 0.1 s, or 18 +/− 1 s and 1.8 s +/− 0.2 s depending on the labeling strategy (Fig. 3D). Considering that elongating ribosomes has been estimated to proceed at a speed of 16-17 amino acids per second on average under similar growth conditions, i.e. 2 doublings/h^34–35^, our estimates for the dwell-time in the slow diffusion state, 15-18 s, are in a good agreement with the time required to translate one ‘typical’ *E. coli* protein with a length of ~250 amino acids^36^ which would take 15-16 s.

The fact that the slow diffusion state dwell-time is the same for both subunits (Fig. 3D), have further interesting consequences. More than 50% of *E. coli* genes are organized in operons which are transcribed into polycistronic transcripts^37^. Polycistronic organization of the transcriptome was shown to be important for translational coupling of genes for which the exact stoichiometry is required for correct operation^38–40^. Such coupling has been suggested to occur through either or both of two hypothetical mechanisms^41^. In the first case, translation of a downstream gene by a ribosome only occurs when another ribosome is translating the upstream gene, thereby unwinding a stable mRNA structure, which would otherwise block the ribosome binding site of the downstream gene^41^. In the second scenario, initiation on the downstream gene occurs by the very same 30S^42^ or 70S^43^ which translated the upstream gene, without dissociation from the mRNA. Our estimates for the dwell-time in the mRNA bound diffusion state of both the small and the large ribosomal subunit (Fig. 3D), strongly suggest that, in the majority of mRNA binding events, ribosomes translate only *one gene at a time*. Further, since we see no apparent difference in dwell-times between the small and the large subunits (Fig. 3D), independent of model size (Fig. 3B), *both subunits are likely to dissociate from an mRNA after translation termination*.

### Selection of initiation sites by 30S subunits

During translation initiation, the 30S subunit is required to rapidly scan the transcriptome in order to identify a correct start codon in a vast pool of potential sites. In eukaryotes, the identification of initiation sites on mRNA by the small ribosomal subunit is not direct, but requires initiation factors, specific mRNA structures, and mRNA scanning^44^. In *E. coli*, however, initiation sites on mRNAs can be recognized by 30S alone^45^, and since the majority of bacterial mRNAs are polycistronic, initiation sites should be searched throughout the entire transcripts, and not only by the 5’ end. In certain cases, diffusion along an mRNA has been shown to promote translation initiation^42^, but it is generally assumed that the 30S subunit binds directly or in very close proximity to the initiation site. Base pairing between SD and ASD should, in principle, direct the 30S to the start codon. However, a recent study employing ribosomes with a modified ASD sequence, provided strong evidence that the SD-ASD interaction is far from enough for efficient translation, and that initiation sites are rather hardwired by other mRNA features^28^. It has also been shown that the initial association of 30S with mRNAs depends on secondary structures by the initiation site, independent of the SD motif and start codon, which are monitored in subsequent steps^46–47^. Yet, it is still debated to what extent initiation relies on SD-ASD interaction in *E. coli* and how widespread this mechanism of modulation of translation initiation is in other bacteria^48–49^.

In the above experiments, which helped us to establish the threshold for separation of free and mRNA-bound 30S subunits, we noticed that the previously developed O-30S^25^ appear to be significantly involved in binding of endogenous mRNAs (Fig. 2A-B). Besides a mutated ASD sequence, these ribosomes carry the additional 16S rRNA substitutions G722C and U723A located in the vicinity of the helix formed by SD and ASD, as well as a C1192U mutation which confers spectinomycin resistance^50^. To investigate the contribution from the ASD sequence alone, as well as these additional rRNA mutations, we analyzed the occupancy and dwell-times of several modified 30S subunits in the freely diffusing state and the mRNA-bound state using the coarse-grained HMM-analysis as described above (Tables S15-S18). To minimize the effect of toxicity associated with expression of ASD-modified ribosomes, these experiments were performed with new constructs, with only low-level expression of the mutated rRNA operons.

For 30S subunits in which only the ASD sequence was mutated (O-ASD), a substantial fraction (69%) was still found to be in the mRNA-bound state at any given moment, although the corresponding O-SD is not present in the cells. Incorporation of the G722C-U723A mutations (O-ASD-C722-A723) and the additional C1192U mutation (O-30S) decrease this fraction further, to 57% and 53% respectively, whereas the additional C1400U mutation (O-30S-U1400) finally brings the bound-state occupancy down to a mere 23% (Fig. 4A).

**Fig. 4.**
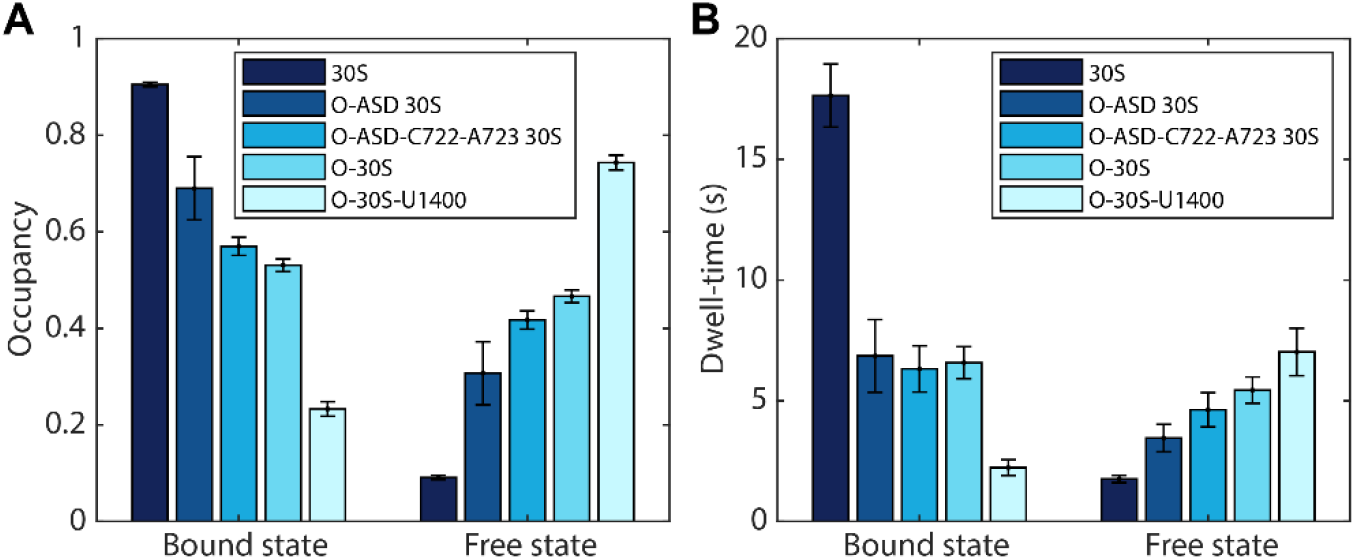
Translation kinetics of modified 30S subunits. Estimated occupancy (**A**) and dwell-time (**B**) of tracked 30S subunit variants in the mRNA-bound state or the freely diffusive state. The data shows the coarse-grained results from HMM-fitted model size 6. Results for all model sizes (2-8), are shown in Table S6 and Tables S15-S18. Error bars represent bootstrap estimates of standard errors.

From the HMM estimated transition frequencies between the free and mRNA-bound diffusion states, we find that O-ASD, O-ASD-C722-A723, and O-30S spend, on average, approximately the same 6.5 s in the mRNA-bound state, whereas the time spent freely diffusing between binding events is between 3.5 s and 5.5 s (Fig. 4B). The relatively long-lived bound state for these mutants suggests their involvement in translation, although it appears to be significantly shorter than what we have calculated for unmodified ribosomes (16-18 s, Fig. 3D and 4B). Selective ribosome profiling on ribosomes with altered ASD sequences has revealed vast differences in translation efficiencies for different genes between wt ribosomes and ASD mutant ribosomes^28^. Based on published data of the relative ribosomal occupancies on individual genes for the O-ASD mutant^28^, we estimate the average length of polypeptides translated by O-ASD ribosomes to be ~150 amino acids (see Methods). Indeed, this is much shorter than the ‘average’ protein translated by wt ribosomes, ~250 amino acids^36^. Moreover, close investigation of ribosome density profiles for individual genes revealed that in a substantial number of instances, initiation occurs at alternative start codons in which orthogonal ribosomes are likely to translate only short nonsense ORFs. Hence, we conclude that the observed binding events in O-ASD-mutant tracking likely represent active translation, albeit of, on average, shorter sequences.

On average it takes only twice longer for O-ASD ribosomes to find initiation sites relative to unmodified 30S (3.5 s +/− 0.6 s vs 1.8 s +/− 0.2 s, Fig. 4B). Incorporation of mutations G722C-U723A and C1192U lead to slightly longer time needed for initiation (4.6 s +/− 0.7 s and 5.4 s +/− 0.5 s, respectively), probably reflecting less affinity to mRNAs. These three 30S variants, however, are likely to have similar preferences in what genes to translate, given the similar average dwell-times in the bound state (Fig. 4B). Finally, the most severely affected 30S mutant, with the additional C1400U mutation in the decoding center, shows dramatically different transition frequencies between free and mRNA-bound diffusion states, with a long-lived free state (7 s +/− 1 s), interrupted only by relatively short binding events for 2.2 s +/− 0.3 s (Fig. 4B). Hence, these orthogonal subunits seem not to translate endogenous mRNAs to basically any extent at all.

## CONCLUSIONS

We have presented a comprehensive analysis of diffusional behavior of *E. coli* ribosomes and developed a toolset to study global kinetics of mRNA translation directly in living cells. Using this approach, we found that both *E. coli* ribosomal subunits are likely to dissociate from the mRNA after translation termination. Hence, we see no evidence of widespread re-initiation on downstream ORFs by 30S^42^ or 70S^43^ remaining bound to the mRNA.

We further examined the importance of the SD-ASD interaction for translation initiation, and show that it has only a modest impact on the overall initiation efficiency. Rather, other features of mRNAs play major roles for regulation of protein expression. It will, thus, be interesting, to further investigate the co-evolution of the transcriptome with ribosomes, and look for specific features of the ribosomes which enable specific interactions with mRNAs. Such findings would significantly help in the development of artificial ribosomes with higher level of orthogonality for bioengineering applications.

Finally, our single-molecule tracking method allows quick evaluation of the translational state of cells, and detection of even subtle changes in state occupancies or state transition rates. After preparation of cells with labeled ribosomal subunits, this information is readily accessible within minutes without any downstream processing after data acquisition, thus minimizing the risk of artefacts. The method further allows for testing of multiple conditions and to introduce changes over the course of the experiment. Hence, the method opens up new possibilities to investigate how bacteria rapidly adapt to changing conditions, such as antibiotics treatment, and regulate their protein synthesis machinery for optimal recourse allocation.

## ACKNOWLEDGMENTS

The authors would like to thank the laboratory of Luke Lavis for providing the JF549-HaloTag dye. This work was supported by The Swedish Research Council (M.J. 2015-04111, 2016-06264, 2019-03714; J.E. and M.J. 2016-06213), the Knut and Alice Wallenberg Foundation (J.E.), and the Carl Trygger Foundation (M.J. 15:243).

## METHODS

### Construction of bacterial strains and growth conditions

For cloning purposes all PCRs were done using Q5 high-fidelity DNA polymerase (NEB) and Gibson Assembly was performed using NEBuilder HiFi DNA assembly master mix (NEB) according to manufacturer’s protocols. Strains expressing L1-, L9-, L25, and S2-HaloTag were constructed using λ Red assisted recombineering^51^. First, the pKD4 plasmid (Addgene #45605) was amplified by inverse PCR using primers pKD4_F and pKD4_R (Table 1) and Gibson-assembled with the HaloTag gene amplified with primers HaloTag_F and HaloTag_R (Table 1) and pFA6a-HaloTag-KanMX6 (Addgene #87029) as a template. Next, the HaloTag-KmR containing fragments were amplified from the resulted pKD4-HaloTag plasmid with pairs of primers X-Halotag-F and X-Halotag-R, where X is the ribosomal protein to be mutated (Table 1). The amplified fragments were used for recombineering to insert HaloTag-KmR cassettes in the genome of *E. coli* MG1655 at the C-terminus of each selected gene^51^.

**Table 1.**
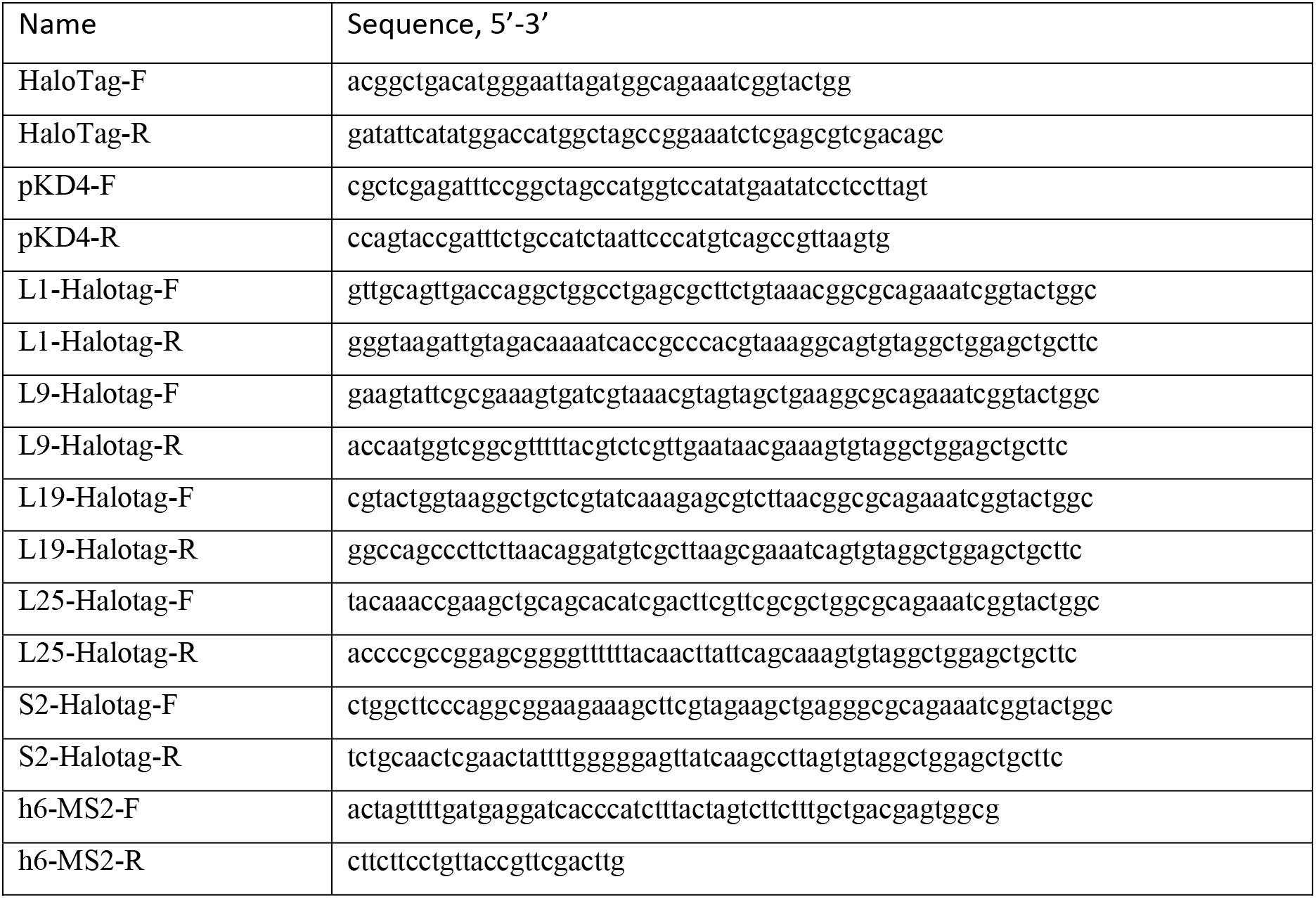
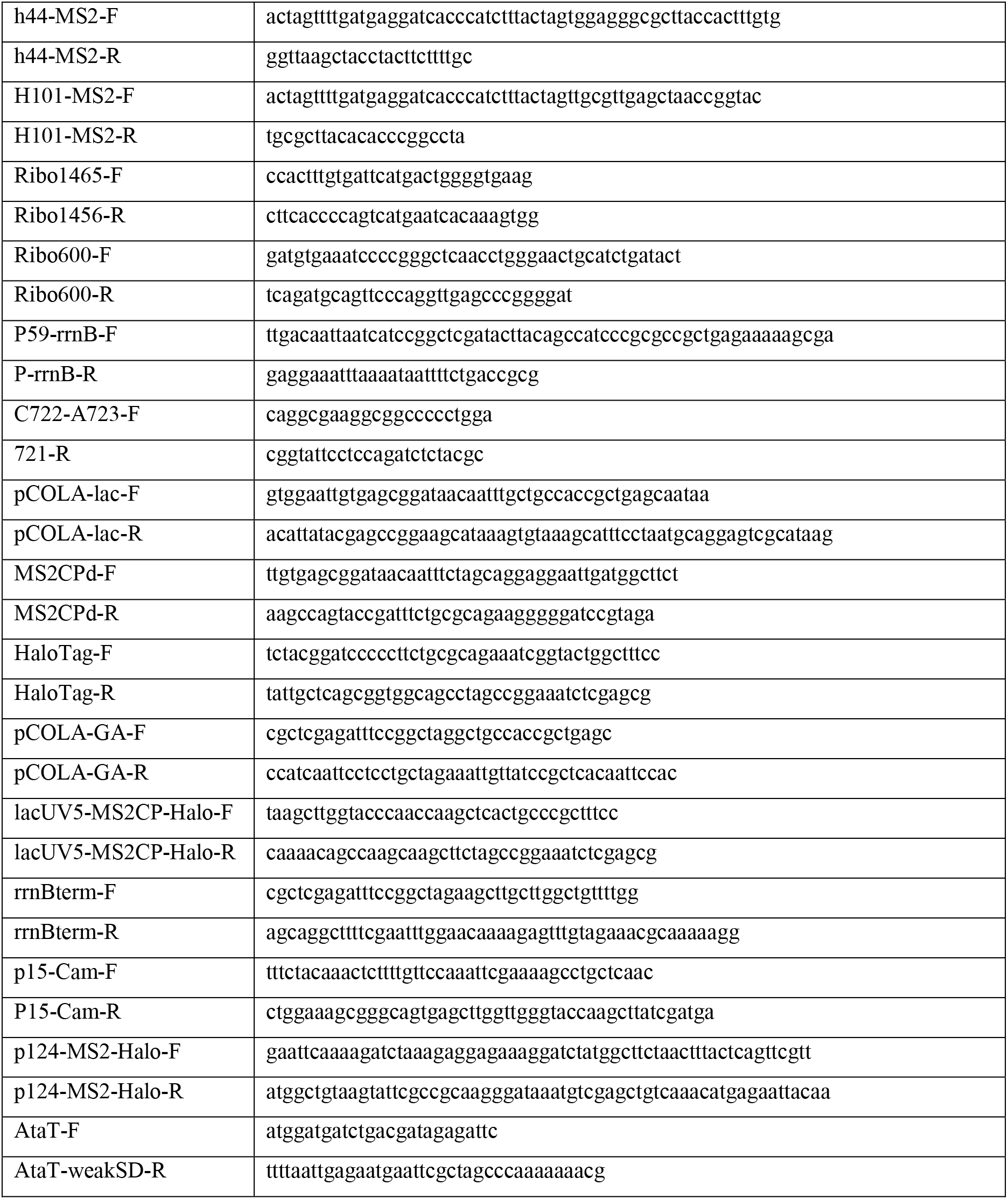
Oligonucleotides used for cloning.

Insertion of the MS2 aptamer into different positions of the *rrnB* operon encoded in the pAM552 plasmid (a gift from A. Mankin lab, as in Addgene #154131) was performed by inverse PCR using primers Y-MS2-F and Y-MS2-R (Table 1), where Y designate the ribosomal helix which was modified, followed by re-circularization through PNK phosphorylation of PCR product ends and ligation by T4 ligase (NEB). Similarly, to create a plasmid for tracking of ribosomes with mutated ASD sequence, the MS2 aptamer insertion at h6 was performed by inverse PCR of the pSC101-O-ribosome (a gift from J. Chin lab) using primers h6-MS2-F and h6-MS2-R (Table 1) and subsequent re-circularization resulting in plasmid pSC101-O-ribosome-h6-MS2. Selection of mutation C1400U in 16 rRNA in pSC101-O-ribosome-h6-MS2 was confirmed by Sanger sequencing of isolated plasmids from independent colonies showing faster growth on LB plates.

To create a plasmid in which only the ASD sequence is mutated and additional mutations (G722C, U723A, and C1192U) are not present, the fragment containing the corresponding positions was amplified using oligonucleotides Ribo600-F and Ribo1465-R (Table 1) from the pAM552 plasmid, and the PCR product was Gibson assembled with the fragment of the plasmid pSC101-O-ribosome-h6-MS2 amplified using primers Ribo1465-F and Ribo600-R (Table 1). The resulting plasmid pSC101-oASD-h6-MS2, when transformed in *E. coli* MG1655, caused high level of toxicity and instability. Therefore, a set of plasmids with a weaker apFAB59 promoter from the BIOFAB collection^52^ (p59) regulating expression of mutated ribosomal operon was created. First, the P1-P2 promoter present in pSC101-O-ribosome-h6-MS2 was exchanged with the p59 promoter by inverse PCR using primers P59-rrnB-F and P-rrnB-R (Table 1), and subsequent re-circularization resulting in plasmid pSC101-P59-O-ribosome-h6-MS2. Next, a derivative of this plasmid in which only the ASD sequence is mutated was created using Gibson Assembly as explained above, yielding the plasmid pSC101-P59-oASD-h6-MS2. Finally, the plasmid pSC101-P59-oASD-C722-A723-h6-MS2, in which 16S rRNA contains the mutated ASD sequence and substitutions G722C and U723A, was created by inverse PCR using pSC101-P59-oASD-h6-MS2 as a template and primers C722-A723-F and 721-R (Table 1), with subsequent re-circularization.

Several plasmids that allow expression of MS2CP-HaloTag were created. First, the lacUV5 promoter was inserted in the pCOLADuet vector (Novagen) by inverse PCR using primers pCOLA-lac-F and pCOLA-lac-R (Table 1), followed by re-circularization by ligation which resulted in the pCOLA-lacUV5 plasmid. Then, the gene for single-chain tandem dimer of the high affinity version of MS2 coat protein and the gene for HaloTag were Gibson-assembled with the pCOLA-lacUV5 plasmid. Corresponding fragments were obtained by PCR amplification using pairs of primers: i) MS2CPd-F and MS2CPd-R for amplification of MS2CP; ii) HaloTag-F and HaloTag-R for amplification of HaloTag; iii) pCOLA-GA-F and pCOLA-GA-R for the linearized plasmid (Table 1). This resulted in in the pCOLA-lacUV5-MS2CP-HaloTag plasmid. Next, a fragment from the pCOLA-lacUV5-MS2CP-HaloTag plasmid, containing the *lacI* gene, lacUV5 promoter and MS2CP-HaloTag fusion, was amplified with primers MS2CP-Halo-F (Table 1) and MS2CP-Halo-R (Table 1). This fragment was fused via Gibson Assembly with two fragments: i) a DNA fragment containing the p15 origin of replication and the gene for chloramphenicol acetyltransferase, and ii) the *rrnB* terminator sequence. The first fragment was amplified using p15-Cam-F and p15-Cam-R (Table 1) from the plasmid pEVOL-pBpF (Addgene #31190), and the second fragment was obtained from the pBAD/HisB vector (Invitrogen) by amplification using primers rrnBterm-F and rrnBterm-R (Table 1). The resulting plasmid is called p15-lacUV5-MS2CP-HaloTag. Finally, a derivative of this plasmid in which the *lacI* gene was removed and the lacUV5 promoter was swapped with a weak constitutive promoter apFAB124 from the BIOFAB collection^52^ was created by inverse PCR using primers p124-MS2-Halo-F and p124-MS2-Halo-R (Table 1) and subsequent re-circularization.

A plasmid pBAD30-ataT containing the gene for the AtaT toxin (a gift from S. Dubiley) from the AtaRT toxin-antitoxin system with a *P*_*BAD*_ promoter regulating the expression^31^ was modified to reduce the expression level by substituting the strong AGGAGG Shine-Dalgarno to the weak TTCTCA sequence by inverse PCR using primers AtaT-F and AtaT-weakSD-R (Table 1) and subsequent re-circularization resulting in plasmid pBAD30-SD_weak_-AtaT.

### Calculation of doubling time

Bacterial strains were grown on LB agar plates overnight at 37 °C. For each tested strain, six individual colonies were grown in LB media until they reached OD_600_=0.1-1. The cells were then diluted to OD_600_=0.0001 in fresh LB and growth kinetics were recorded at 37 °C in a 96-well microplate (100 μl/well) using Spark 10M microplate reader (Tecan) with measurements of OD_600_ performed every 5 min after 1 min shaking. After subtraction of medium background, the minimal doubling time was calculated using a custom MATLAB script, which fits growth curves of cells with an exponential function and identifies 30 min intervals of maximal growth rate for each curve. The data from six independent wells were fitted separately and the results were averaged.

### HaloTag labeling and sample preparation

*E. coli* cells containing HaloTag were grown on LB agar plates overnight at 37 °C. Several colonies were used to inoculate EZ Rich Defined Medium (RDM, Teknova) supplemented with 0.2% glucose. Cells were grown at 37 °C with shaking to OD=0.5-1, harvested by centrifugation at 2000 g, resuspended in 150 μl of fresh RDM media, and JF549 HaloTag ligand (a gift from Lavis lab) was added to the final concentration of 0.5 μM. After incubation with the ligand for 30 min at 25 °C, cells were washed twice using M9 media supplemented with 0.2% glucose, then incubated in RDM media at 37 °C with shaking for 60 min to facilitate the release of unbound ligand from cells. After incubation, cells were washed 3 times using M9 media supplemented with 0.2% glucose. Finally, cells were resuspended in RDM to OD_600_ ≈ 0.003 and sparsely spread onto an agarose pad prepared with RDM and 2% agarose (SeaPlaque GTG Agarose, Lonza). The sample was placed on the microscope with a cage incubator maintaining temperature at 37 ± 2 °C. Cells were grown for 2 hours forming mini-colonies before image acquisition.

For KSG treatment experiments, cells were incubated on agarose pads at 37 °C for 2 hours after which RDM media supplemented with 0.2% glucose and containing either 50 μg/ml or 500 μg/ml KSG was injected to the sample with growing mini-colonies. Imaging of cells was performed 60 min after injection of antibiotic. For experiment in which production of AtaT toxin was induced, *E. coli* MG1655 expressing S2- or L9-HaloTag and transformed with the pBAD30-SD_weak_-AtaT plasmid were grown on an agarose pad to form mini-colonies for 2 hours at 37 °C. Production of the toxin was induced by injection of RDM media supplemented with 0.2% glucose and 0.2% arabinose. Imaging was performed 70 min later.

Each experimental dataset included 2–5 replicated microscopy experiments, each comprising 30–100 cell colonies (8-100 cells each). The data were found to be consistent in between different repetitions and were combined for analysis.

### Optical Setup

An inverted microscope (Nikon Ti-E) in combination with a CFI Apo TIRF 100 × 1.49 NA objective (Nikon) was used. Bright-field and fluorescence images were acquired using an iXon 897 Ultra EMCCD camera (Andor) coupled to an additional 2.0 × lens (Diagnostic Instruments DD20NLT). Phase contrast imaging was performed with an Infinity2-5 M camera (Lumenera). To track JF549 bound to HaloTag, a 553 nm laser (SLIM-553, Oxxius, 150mW) with a power density of 3 kW/cm^2^ on the sample plane was used in stroboscopic illumination mode with 3 ms laser pulses per 30 ms camera exposures. The microscope was controlled using the μManager software package. Data acquisition from multiple positions was performed using custom μManager plugins.

### Single-particle tracking and analysis of trajectories

Data analysis was performed using custom-made analysis pipelines written in MATLAB (available upon request). Single particle tracking was performed as described previously^6^. Briefly, boundaries of individual cells were extracted from phase contrast images using a previously developed algorithm^53^. Incorrectly segmented cells were manually removed and discarded from the subsequent analysis. Fluorescent spots were detected using the radial symmetry-based method^54^. Refinement of spot positions and estimation of position uncertainties was performed using a symmetric Gaussian spot modelling and the maximum aposteriori fitting^20^. Two-dimensional trajectory building was done using uTrack^55^. Trajectories were built in cells starting from the time-point when only one spot remained in the cell in current and following frames, allowing gaps with 1 missing point. Spots with width (std) > 280 nm, with amplitute < 50 photons, or outside live cells (> 3 pixels outside of boundaries) were excluded. Using a previously described HMM algorithm^6, 20^ we analyzed diffusional properties of tracked molecules by fitting all trajectories with length of ≥ 20 steps obtained for each experimental condition to a pre-set number of diffusion states. The algorithm makes explicit use of spot coordinates as well as localization uncertainties of each spot and handles missing data points^20^. Mean dwell times were calculated from the diagonal elements of the resulted HMM transition matrix, i.e., from the average probability of exiting a diffusive state, thus allowing the estimation of dwell times which are longer than the span of trajectories. To condense HMM models with many states to a coarse grained three-state model, the hidden states were classified as “mRNA bound”, “free”, and “unphysiological” using threshold values of 0.25 μm^2^/s and 1.5 μm^2^/s.

For mean square displacement (MSD) analysis, trajectories were sliced into segments of 7 positions, and linear fits passing through the origin was made using the equation *MSD(t)* = 2*nD*Δt ∙ t, where *t* = 1, 2, 3 (first 3 points), *n* = 2 is the dimension of the trajectories, Δ*t* = 30*ms* is the time step, and the apparent diffusion coefficient *D* is the fitting parameter.

### Analysis of previously published ribosome profiling data

The average length of polypeptides synthesized by O-ASD 30S was calculated from the ribosome density fraction on the top hundred most translated genes (comprising about 2/3 of all translation) in the O-ASD 30S ribosome profiling data in^28^.

## SUPPLEMENTARY MATERIAL

**Supplementary Movie S1.** Fluorescence microscopy data of L9-HaloTag labeled 50S ribosomal subunits, acquired with 3 ms laser exposures per 30 ms camera frame. Playback is 33 frames per second (i.e. real-time). The left panel shows raw data, whereas the right panel shows the cell outlines (segmented from phase-contrast images) as well as automatically detected diffusion trajectories, color-coded with respect to coarse-grained diffusion states estimated from HMM analysis. Diffusion trajectories were built in cells from the time-point where only one spot per cell remained in the current and all subsequent frames.

**Fig. S1.**
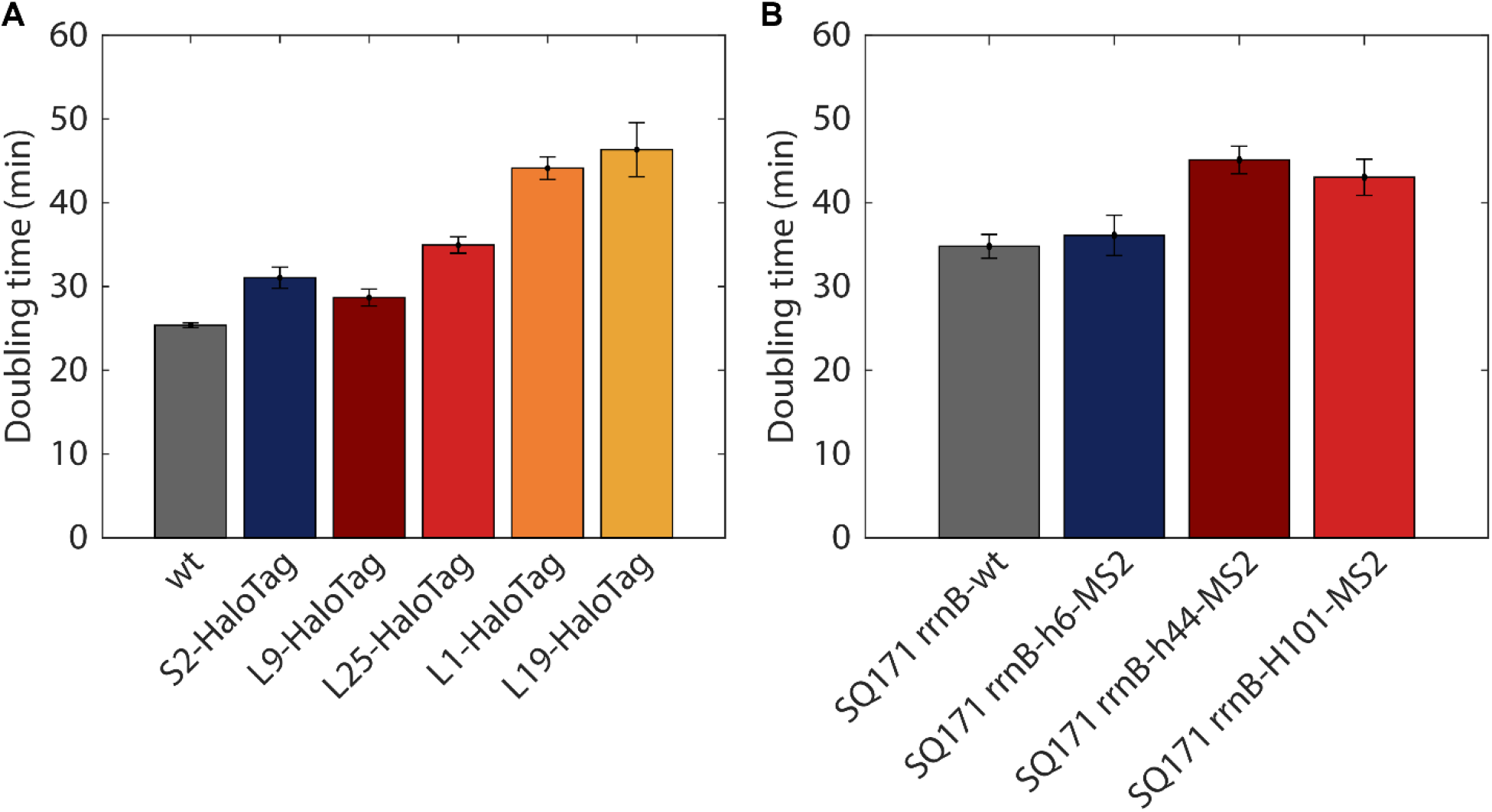
Growth of *E. coli* strains constructed for tracking of ribosomal subunits. **A**. Doubling times of *E. coli* cells expressing S2-HaloTag, L1-HaloTag, L9-HaloTag, L19-HaloTag, L25-HaloTag, and isogenic *E. coli* MG1655. Error bars represent standard deviation calculated from 6 independent replicates. **B**. Doubling time of *E. coli SQ171* strain which lacks all chromosomal rRNA operons and carrying plasmid pAMM552 expressing either the wt *rrnB* operon or its mutated versions in which the MS2 aptamer was inserted in one of the helixes (helix 6, helix 44, or H101). Error bars represent standard deviation calculated from 6 independent replicates.

**Fig. S2.**
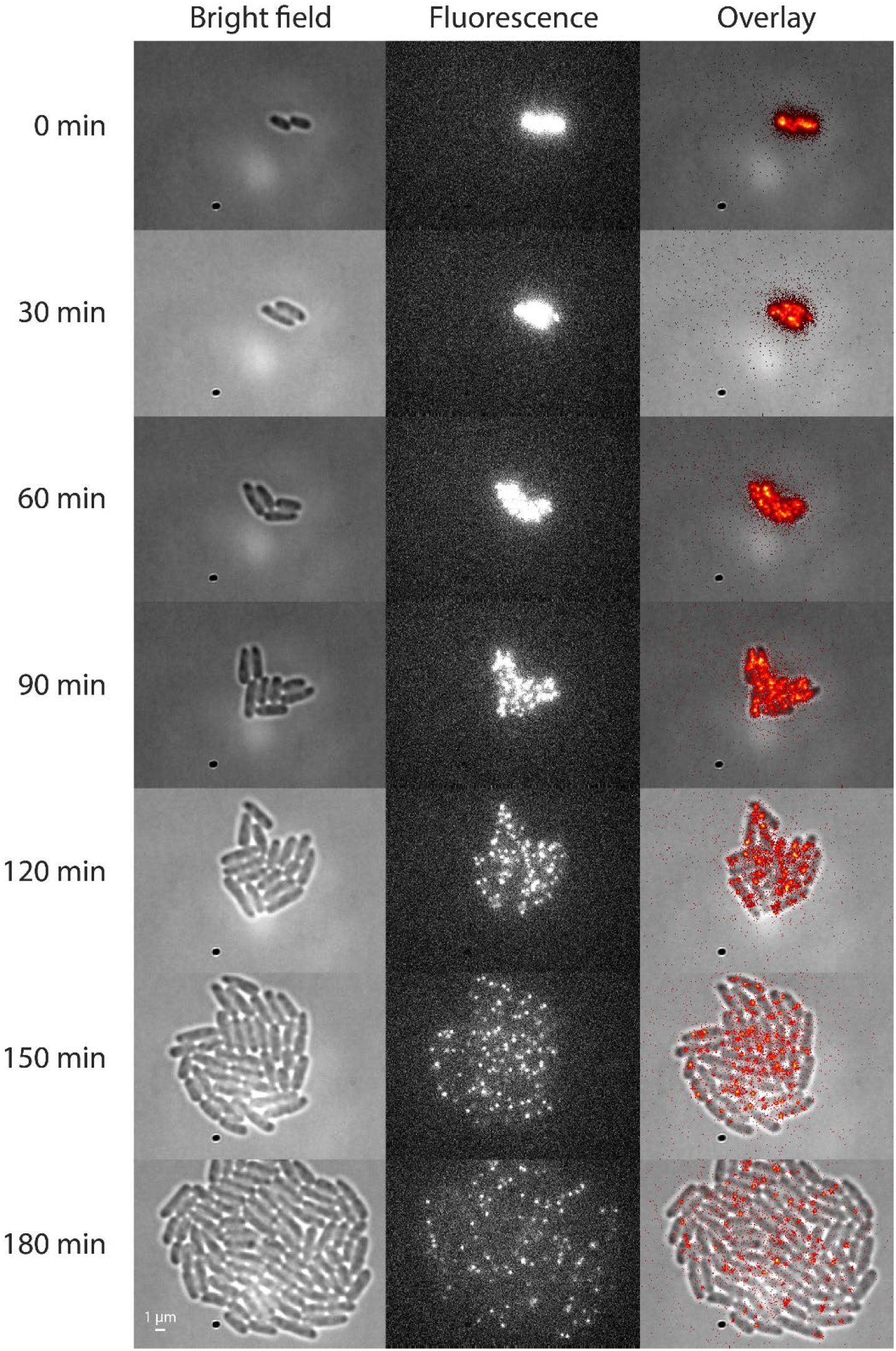
Bright-field and fluorescence images (553 nm laser illumination) of L9-HaloTag expressing *E. coli* cells after labeling with the JF549 HaloTag ligand.

**Fig. S3.**
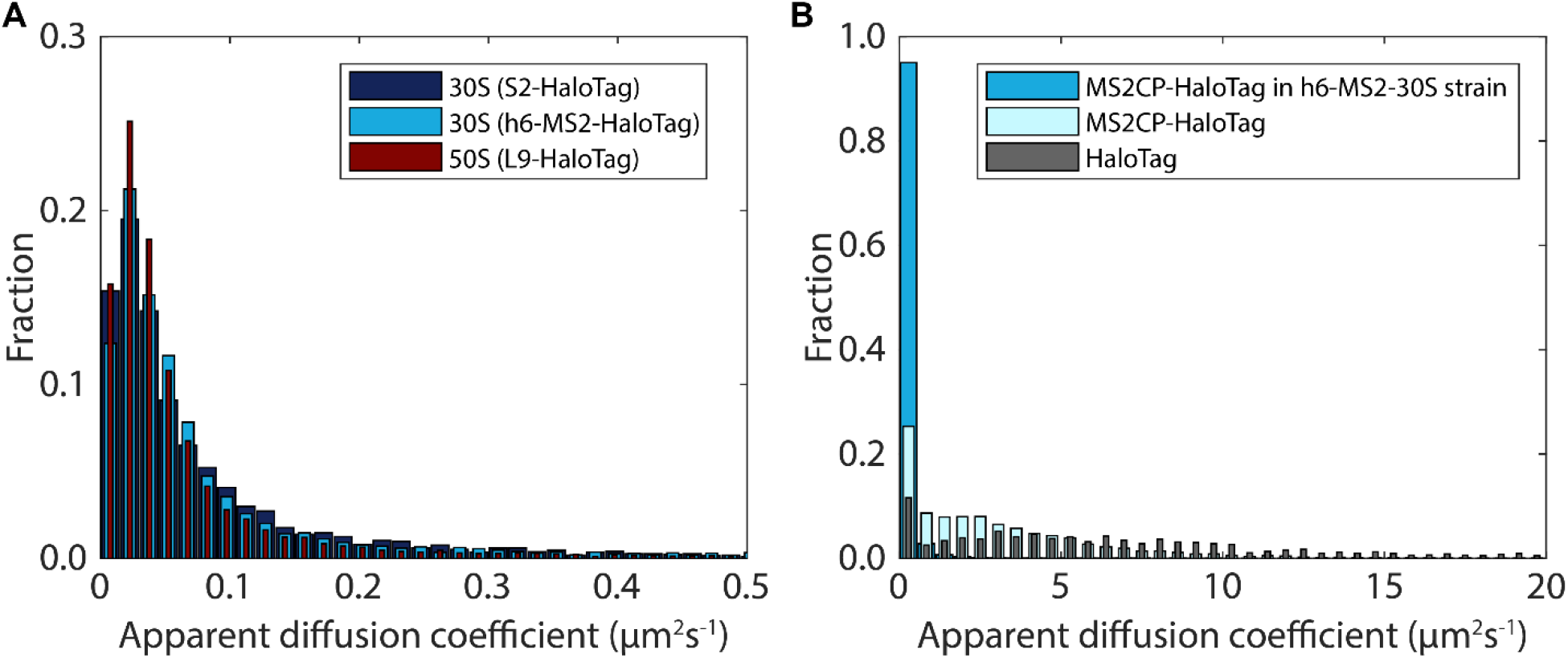
Apparent diffusion coefficients estimated from mean-squared-displacement. **A**. Distribution of apparent diffusion coefficient for 30S or 50S HaloTag labeled ribosomal subunits. **B**. Distribution of apparent diffusion coefficient for free HaloTag and the MS2CP-HaloTag fusion in cells lacking the MS2 RNA aptamer. Data for diffusion of the MS2CP-HaloTag fusion in cells with the MS2 RNA aptamer inserted into a subpopulation of the 16S rRNA is shown for comparison. The average diffusion coefficient for HaloTag and the MS2CP-HaloTag were calculated to be 4 μm^2^/s and 12 μm^2^/s, respectively. The apparent diffusion coefficients in both panels were estimated from mean-squared-displacement analysis of diffusion trajectory segments of 7 frames. Fluorescence data were acquired at 30 ms frames for panel A, and 5 ms for panel B.

**Fig. S4.**
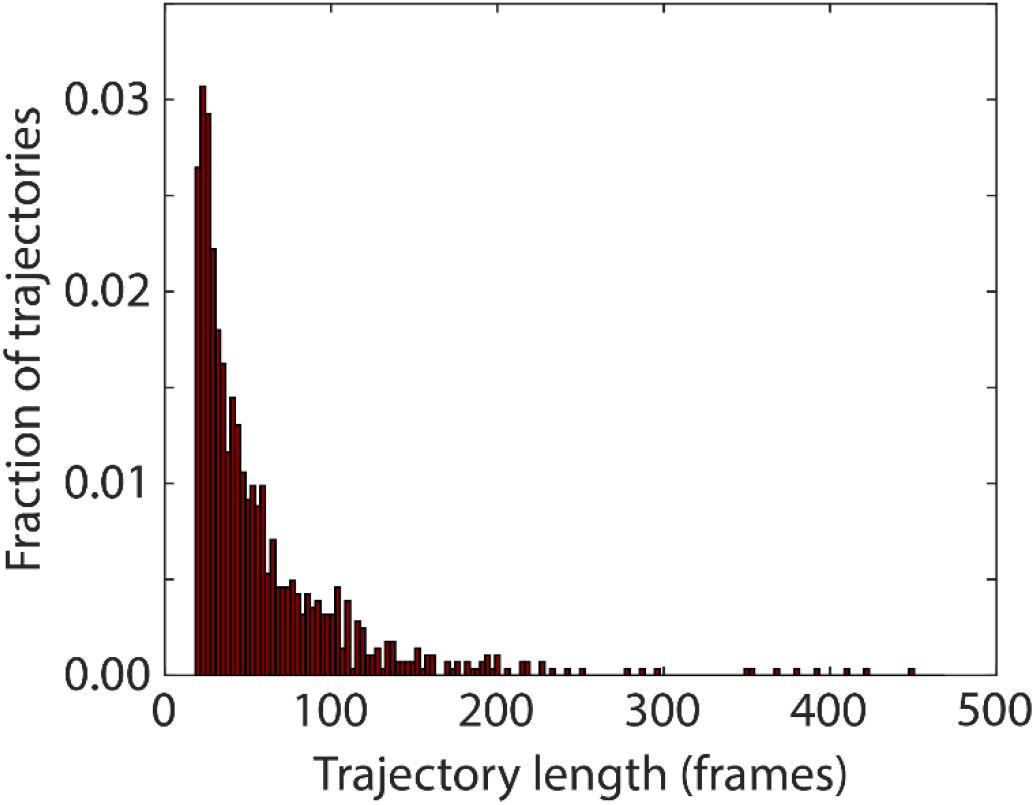
Distribution of trajectory lengths from a single 50S (L9-HaloTag) tracking experiment. Only trajectories of length 20 frames or longer were saved for analysis.

**Fig. S5.**
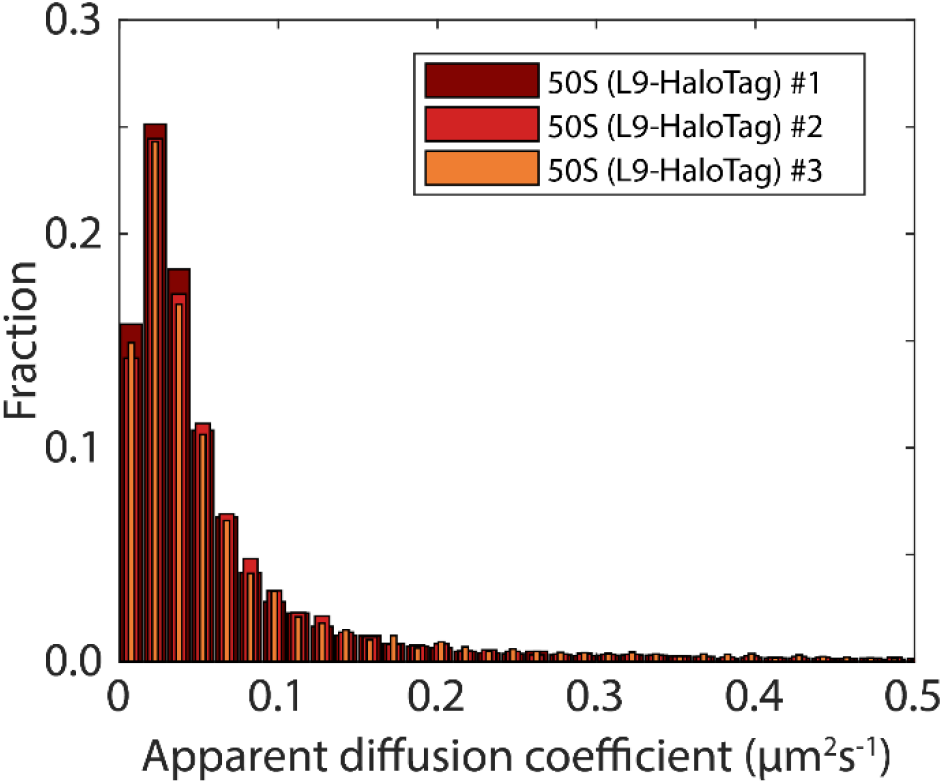
Distribution of apparent diffusion coefficient for L9-HaloTag labeled 50S ribosomal subunits, estimated from mean-squared-displacement analysis of diffusion trajectory segments of 7 frames from three separate experiments.

**Fig. S6.**
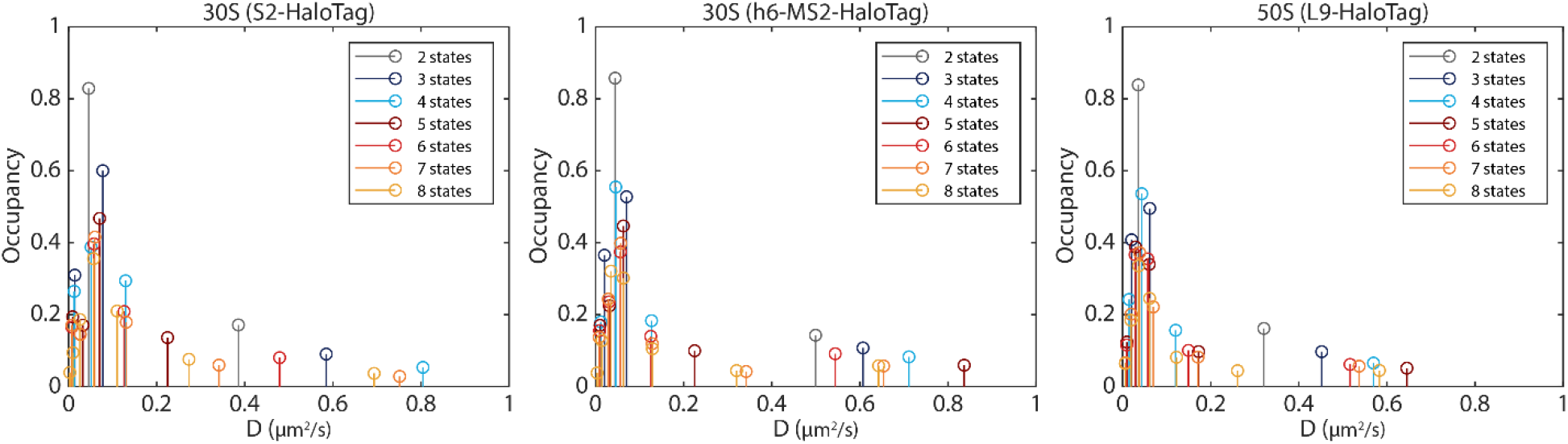
HMM-estimated occupancies in different diffusion states for all fitted model sizes. All datasets, including low-occupancy states at D > 1 μm^2^/s, are also shown in Tables S4-S6.

